# Knowledge “installed” diffusion model predicts the geometry of actin cytoskeleton from cell morphology

**DOI:** 10.1101/2023.01.12.523863

**Authors:** Honghan Li, Shiyou Liu, Shinji Deguchi, Daiki Matsunaga

**Affiliations:** Division of Bioengineering, Graduate School of Engineering Science, Osaka University, Osaka, Japan; School of Life Science, Peking University, Beijing, China

## Abstract

Cells exhibit various morphological characteristics due to their physiological activities, and changes in cell morphology are inherently accompanied by the assembly and disassembly of the actin cytoskeleton. Stress fibers are a prominent component of the actin-based intracellular structure and are highly involved in numerous physiological processes, e.g., mechanotransduction and maintenance of cell morphology. Although it is widely accepted that variations in cell geometry interact with the distribution and localization of stress fibers, it remains unclear if there are underlying geometric principles between the cell morphology and actin cytoskeleton. Here we present a machine learning system, which uses the diffusion model, that can convert the cell shape to the distribution of stress fibers. By training with corresponding datasets of cell shape and stress fibers, our system learns the conversion to generate the stress fiber images from its corresponding cell shape. The predicted stress fiber distribution has good agreement with the experimental data, and the overlap region of predicted and experimentally observed stress fibers reaches 79.3 ±12.4%. We then found some unknown natures such as a linear relation relationship between the stress fiber length and cell area. With this “installed” conversion relation between cellular morphology and corresponding stress fibers’ localization, our system could perform virtual experiments that provide a visual map showing the probability of stress fiber distribution from the virtual cell shape. Our system provides a powerful approach to seek further hidden geometric principles between the cell morphologies and actin cytoskeletons.

## Introduction

Actin stress fibers (actin SFs) are a prominent component of the actin-based intracellular structures and are mainly composed of actin and myosin [1], which generate strong contractile forces and play a vital role in cell motility and morphological changes [2]. SFs exhibit adaptive responses to changes in intracellular and extracellular cues by modulating their assembly and disassembly [3], which allows them to be involved in diverse cellular functions such as the regulation of cell–substrate adhesions [4, 5], cell migration [6], mechanotransduction [7, 8], morphological maintenance [9, 10], senescence [11], mechanical properties [12], and avoidance of proinflammatory signaling and maintenance of cell homeostasis [13, 14].

Cells sense mechanical properties in their surrounding microenvironment [15, 16] and change their geometric morphologies, both internal cytoskeletons and outer shapes, to modulate their functions. Many previous studies reported the geometric relation between the cell outline and the alignment/localizations of SFs. SFs, or actin filaments are known to align along the longest axis of the cell shape (confinement), and this tendency becomes stronger for cells with large aspect ratios [17–20]. The curvature of the cell contour, which can be obtained by fitting the cell outline with circles, also affects the localization of SFs and the contractile force as a consequence; SFs tend to be to be more concentrated where the local curvature is concave [21], and they are thicker at a place where the curvature radius is larger [22]. The curvature was previously used to estimate the contractile force of the cell [23, 24] since the tension of these SFs is in balance with the membrane tension [25], and this local curvature affects various cell functions such as differentiation [26]. Based on these experimental observations, some previous studies attempted to build simple models to predict the geometry and localizations of SFs out of limited information. Using geometric constraints of micro-patterning, previous studies [24, 27–29] have successfully described the localization of SFs inside cells using one- or two-dimensional bio-chemo-mechanical models. More recently, there is an attempt to predict the alignment of the actin cytoskeleton by modeling the cytoskeleton as a nematic liquid crystal [30]. In this model, they coupled the dynamics of cellular contour and the internal topologies, and discussed how these interplays impact the internal/outline geometries of cells.

Although it is required to understand how the cell shape and cytoskeleton crosstalks in order to adapt to their microenvironments, it is still difficult to draw underlying geometric principles between them from the big data of experiments. We present a machine learning-based system that predicts the geometry and localization of SFs from cell shape. To prepare training data, we extract cell shapes from microscope images using image processing methods and segment SFs using a convolutional neural network (CNN) method. Our machine learning system, which is based on the diffusion model [31], is trained to learn the rules to convert the input cell shape images to new images predicting where SFs are likely to be in the corresponding cell shape. We thus show that our system can recover important geometric features of SFs from the cell shapes. We also propose that this system, which has”installed” knowledge from experiments, can be used to perform virtual experiments, generating SFs localization maps from artificial cell images. The virtual experiments give us access to quantitative evaluation of the SF geometry under desired conditions, allowing for searching hidden geometric principles that underlie between the cell shape and SFs.

## Materials and methods

Fig 1A summarizes our machine learning-based system. The process has four stages: (A) culture cells and acquire the cell images, along with the SFs, (B) extract cell outlines using a sequence of image processing methods, (C) segment the SFs using a CNN method, and (D) train the diffusion model using the SFs and cell shape so that the system understands the correlations between them.

**Fig 1.**
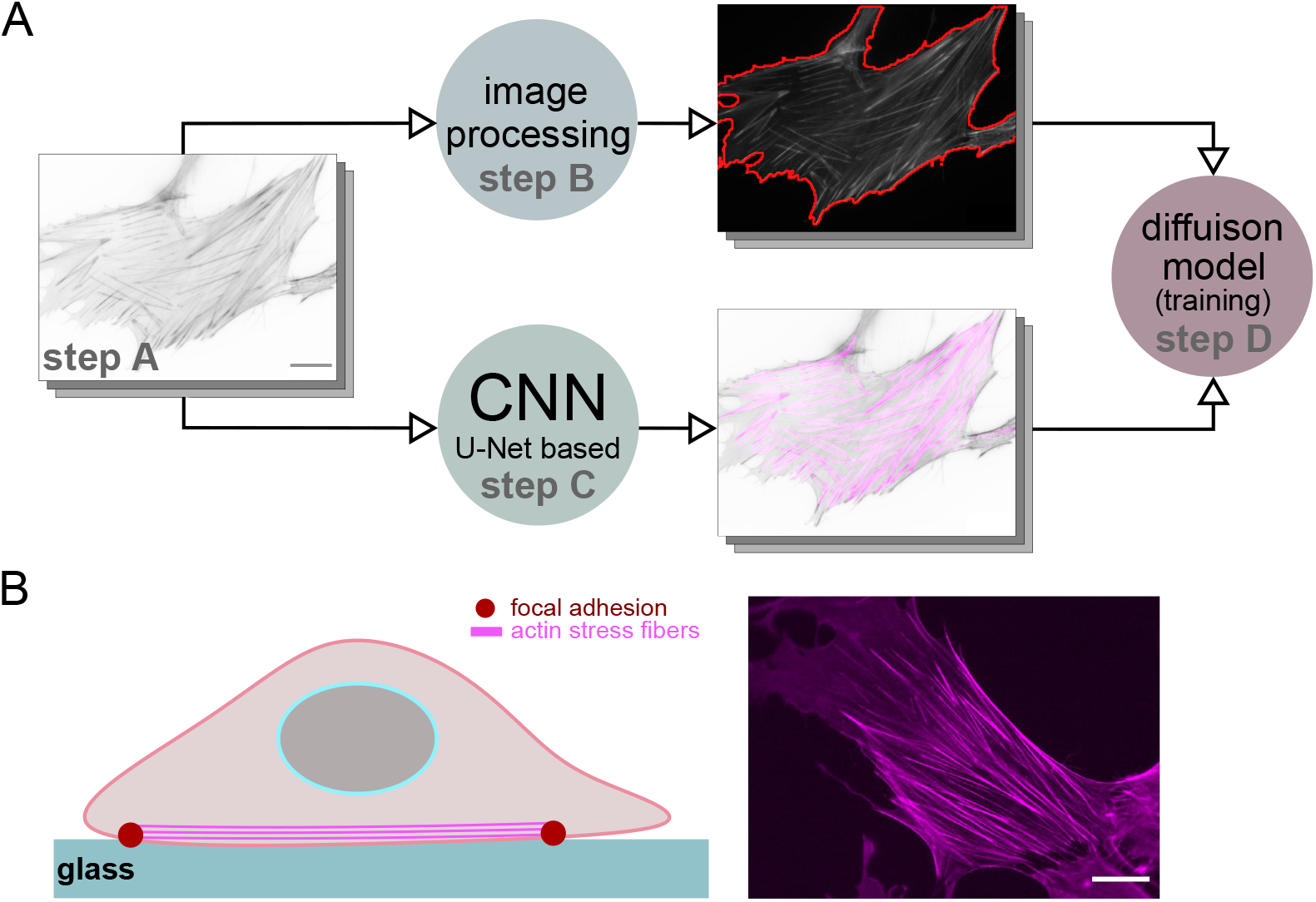
Overview of our approach and experimental material. (A) The base bone of our method. By applying the techniques of image processing and the CNN method, the cell outline and SFs can be extracted from the original microscopic images. The extracted outline and SFs will be sent into the diffusion model so that the network can understand the transformation from cell outlines to SFs. (B) Schematic of cell culture. The cells are cultured on the glass substrate, and the SFs can be visualized by fluorescently-labeled phalloidin. Scale bars present for 20 *µ*m.

### Step A: Experimental material

Culture human foreskin fibroblasts HFF-1 (ATCC) in DMEM with high glucose, l-glutamine, and phenol red (Wako), supplemented with 15% fetal bovine serum (Sigma-Aldrich) and 1% penicillin-streptomycin solution (Wako), in a 5% CO_2_ incubator at 37°C.

Fix cells cultured on glass-bottom dishes with 4% paraformaldehyde in PBS (Wako) at 37°C for 30 min, and permeabilize with 0.1% Triton X-100 for 15 min, as shown in Fig 1B. Stain the actin SFs with fluorescently labeled phalloidin (Thermo Fischer Scientific) to visualize and capture the fluorescence imaging of the cytoskeletons using a confocal laser scanning microscope (FV1000; Olympus) equipped with a UPlan Apo 60× oil objective lens (NA=1.42). The total number of acquired SFs images is 1293.

### Step B: Cell shape extraction

The details of the cell outlines are not sensitive to the image resolution; therefore, all the procedures in the cell outline extraction are executed under the resolution of 256×256, which is downsampled from 1024×1024. The whole process of the image outlines using a sequence of image processing algorithm for the cell outline extraction can be summarized as three basic steps. First, a Gaussian filter is applied to the raw microscopic images to remove the noise and the excessive local gradients caused by SFs, and the kernel size for the Gaussian filter is set to 8. Second, a binarization method with adaptive threshold [32] is applied to separate the cell outline from the background. Third, we applied morphological operations (erosion for two times and dilation two times) to the binarized images, which allowed holes and localized small connected components caused by the binarization to be removed from the images.

### Step C: Actin stress fibers segmentation

We proposed a CNN-based method to segment the SFs from the raw fluorescence microscopic images automatically. First, we prepared 50 original images with a resolution of 1024×1024 for the training data and 10 for the test data. The stress fibers are labeled by manual work, requiring high labor costs and time spent. Therefore, we proposed an image augmentation method to maximize the use of information from the labeled images. Since CNN is usually adept at capturing localized textural information [33], we cut each of the original 50 images into four pieces with equal length and width and combined them randomly, as shown in Fig 2A. During the combination, we utilize several image processing methods on each piece to simulate the data with unpredictable random noise that is introduced in experiments and data acquisition: (i) apply Gaussian filter and mean filter to imitate the out-of-focus state of the microscope, (ii) add noise to simulate the conditions with high background noise, (iii) randomly adjust the abstract and brightness of the images to reduce the impact of exposure intensity for different samples in the experiment during the training, and (iv) apply random clockwise rotation and flipping on the images to increase the diversity of training data.

**Fig 2.**
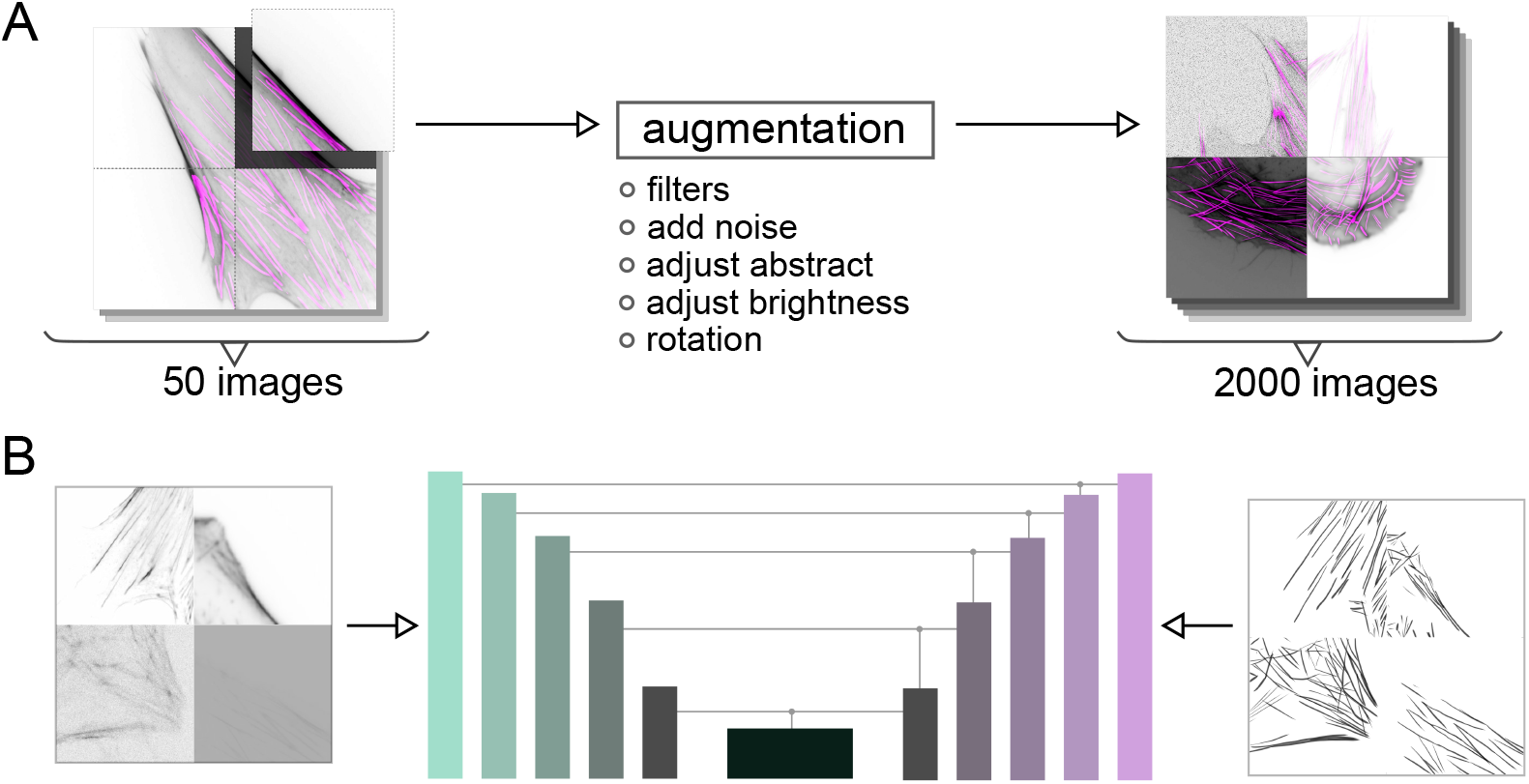
CNN method for the SFs segmentation. (A) Preparation of training data. By utilizing the augmentation methods, the original 50 images are increased to 2000 for the CNN network to be trained. (B) The CNN network (U^2^-Net) is trained with the augmented SFs data. After sufficient training, the network can segment the SFs from microscopic images automatically.

Next, we built a CNN network to segment the SFs, as shown in Fig 2B. The architecture of this network basically refers to U^2^-Net [34], and we deepened the network so that it can process the input augmented SFs data at a resolution of 512 ×512. We train the network by minimizing the KL divergence (binary cross entropy) between the output and ground truth from the training data. We use the training parameters as follows: 50 training epochs (batch size = 4), the parameters *β*_1_ = 0.5 and *β*_2_ = 0.9 are used for the Adam optimizer, *ε* = 2.0 × 10^−4^ learning rate for the optimizer. Nvidia RTX A4000 and Nvidia Titan RTX accelerate the whole training process. After sufficient training, the CNN network could be utilized to segment the SFs.

### Step D: Training with diffusion model

The diffusion model [31, 35] mainly consists of forward diffusion and reverse denoising processes, as shown in Fig 3A. The forward diffusion process can be considered a Markov chain aiming to gradually add Gaussian noise to the input SFs data until it becomes pure Gaussian noise after a total of *T* timesteps. The reverse denoising process is designed to recover the original SFs data from the corrupted data produced by the forward diffusion process. By combining the reverse denoising process with the corresponding cell contour condition, it is possible to remove the noise from the data and recover the original SFs data. This process is also illustrated in Fig 3A.

**Fig 3.**
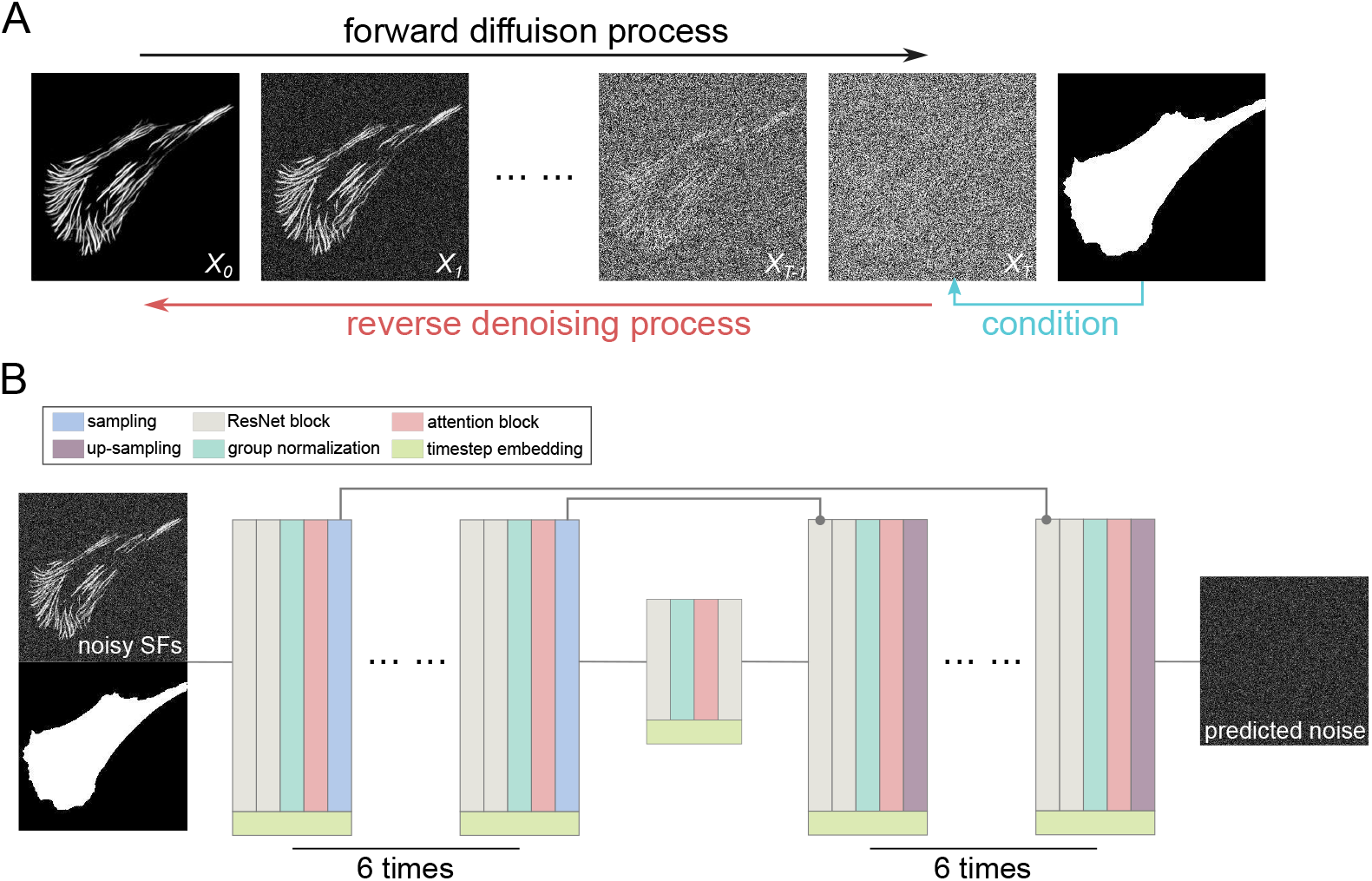
Generating SFs from cell shapes using the diffusion model. (A) Schematic of the diffusion model showing the forward process of turning SFs images into Gaussian noise and the reverse denoising process of transforming Gaussian noise into SFs images under a certain cell contour condition. (B) Architecture of the neural network used to predict noise added to SFs data in the forward diffusion process under the cell contour condition. After training, the network can generate SFs data from random noise based on the input cell contour.

The forward diffusion process [31] can be expressed as follows,

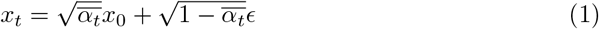

where *x*_0_ are the initial input SFs images, 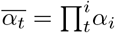 is the variance schedule, and *ϵ* ∼ *N* (0, 1) is the Gaussian noise. Note that *α*_*i*_ is an incremental sequence of values such that 0 *< α*_*T*_ *<* … *< α*_1_ *<* 1. While the reverse denoising noise process aims togradually recover the corrupted *x*_*t*_ to *x*_0_ by estimating the noise in iterations [1, …, *T*].

This process can be modeled [36] as

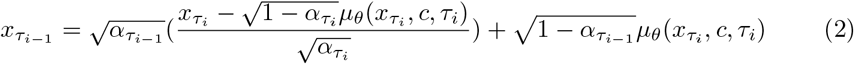

where *τ* is an increasing sub-sequence of [1, …, *T*] with length *S* for accelerating the reverse process [36], *c* denotes cell contour condition and *µ*_*θ*_ denotes the neural network. The object of the neural network is to predict the Gaussian noise *ϵ* that is added in the forward diffusion process based on the input noisy SFs images *x*_*t*_, contour condition *c* and the specific timestep *t*. The object function of this process can be simplified to:

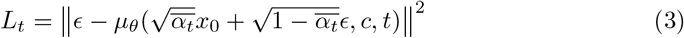

where the neural network tries to minimize *L*_*t*_. The topology of our network is designed refer to U-Net [37] with a contracting path, a bottleneck and an expansive path as shown in Fig 3B. In the contracting path, we designed a module that consists of several process units, which are ResNet block [38] for 2 times, group normalization [39], attention block [40] and sampling operation. While in the expansive path, we use the same module but replace the sampling operation with an up-sampling operation. The bottleneck path has successive modules of ResNet block, group normalization, attention block and ResNet block. Each module in the contracting path and expansive path repeats 6 times, and the output of each module in the contracting path is added to the input of corresponding modules in the expansive path. Each block has an additional concatenation with timestep embedding [41] in order to let the network know the current timestep.

In summary, the whole process of our diffusion model is to train a noise estimation model *µ*_*θ*_ and then generate SFs data from a random noise *ϵ* under the condition of cell shape *c* iteratively. When training the model, the noise *ϵ* is generated randomly first, and this noise is used to destroy the original SFs data *x*_0_. Furthermore the destroyed SFs data are used to predict the noise *µ*_*θ*_(*x*_*t*_, *c, t*), and finally, the predicted noise is expected to be similar to the actual noise *ϵ*. After sufficient training, the network can generate SFs data from random noise according to the input cell shape by gradually subtracting noise based on timesteps. The size of the input Gaussian noise *ϵ* and cell contour images *c* along with the output SFs images *x*_*t*_ are 256×256. The total length of timestep iteration *T* is 800, and the linear method with *τ*_*i*_ = ⌊0.1*i*⌋ is utilized to sample the sub-sequence and the length is *S* = 80. For the variance schedule method, we use the cosine schedule [42] to control 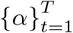. The total number of SFs data is 1293, we randomly choose 1193 (1200 for the last round) images 13 times without duplication for training and 100 (93 for the last round) for the test, and the training images will be randomly rotated and flipped during the training in order to increase the robustness of this machine learning system. We use the training parameters as follows: 1500 training epochs (batch size = 16) and learning rate for the Adam optimizer *ε* = 1.5 × 10^*−*4^.Nvidia RTX A4000 and Nvidia Titan RTX accelerate the whole learning process.

## Results

### Accuracy of the SFs segmentation using CNN

We first evaluate the segmentation performance using CNN. The ground-truth data (*N*_test_ = 10) are produced by manually tracing the stress fibers with straight lines, and we compared our CNN method in Fig 4 with a filament sensor [43], which is an image analysis based method. The comparison is under the resolution of 1344×1024.

**Fig 4.**
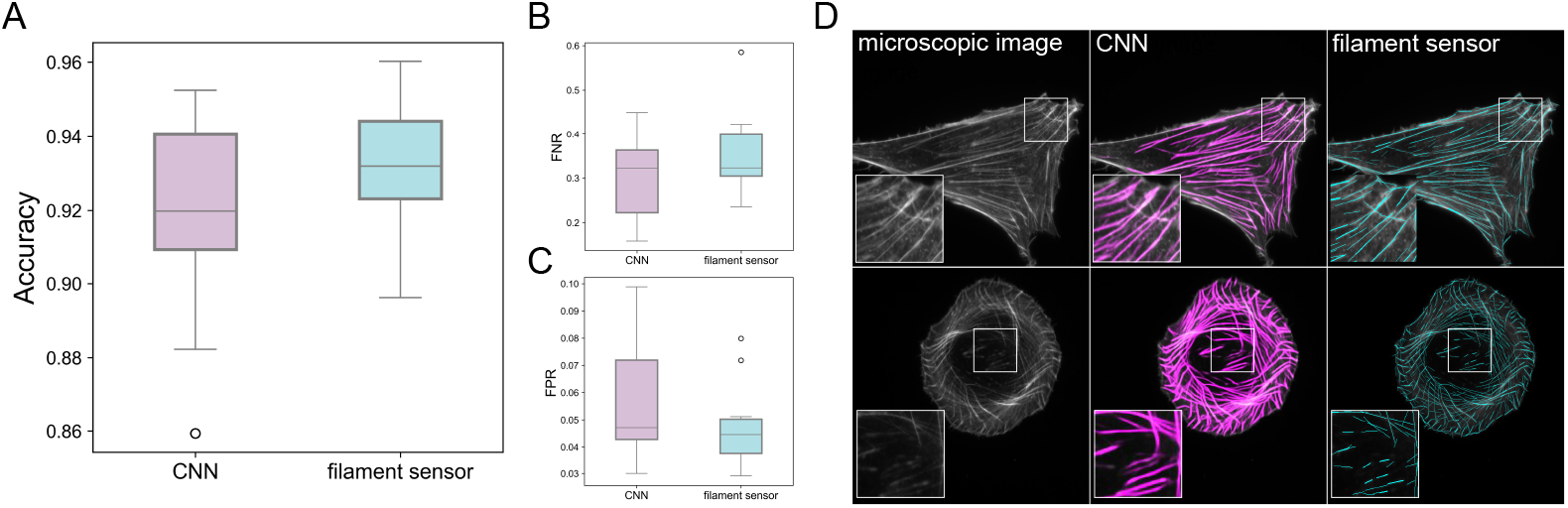
SFs segmentation accuracy using CNN method. (A) Accuracy comparison of SFs segmentation results using our CNN method and the filament sensor from [43]. (B) False negative rate (FNR) and (C) False positive rate (FPR) of SFs segmentation results using our CNN method and the filament sensor, with lower values indicating better performance. (D) Visualization of SFs segmentation results using our CNN method and the filament sensor.

The accuracy of our CNN method and the filament sensor is shown in Fig 4A and D. The accuracy is calculated as the ratio of true positive (TP) and true negative (TN) pixels to the total number of pixels, using a mask size of 8 × 8 to determine the position of SFs. The results show that our CNN method has a similar performance (0.917 ± 0.027) compared to the filament sensor (0.932 ± 0.017). For further comparison, we also compute the false negative rate (FNR) and false positive rate (FPR) as follows: FNR = FN/(TP + FN) and FPR = FP/(FP + TN), where FP and FN represent the number of pixels of false positive and false negative, respectively. Fig 4B and Fig 4C show that our CNN method has a lower FNR (0.302 ± 0.088) and higher FPR (0.055 ± 0.021) compared to the filament sensor (0.351 ± 0.096) and (0.047 ± 0.016), respectively. This indicates that the filament sensor is better at capturing fine details in local SFs, but tends to over-segment them and break the segmented lines at places where the local curvature changes drastically, as seen in Fig 4D.

In conclusion, the filament sensor achieves accuracy that is comparable to machine learning just with image processing techniques. One small negative aspect of this method is that it requires artificial adjustment of hyper-parameters during the pre-processes and binarization processes, which could lead to time-consuming parameter tuning. Although these two methods have similar performance, we choose to use our image augmentation method and the CNN approach, and we now can use these segmented SFs images to train our diffusion model.

### Geometric relation between SFs and cell morphology

This section focuses on how the cell morphology (area, principal direction, aspect ratio, circularity) interacts with the geometry of the actin SFs (total length, principal direction, alignment). Thanks to the automated segmentation method introduced in the previous section, we can now analyze the SFs dynamics under big data (*N* = 1293) with the help of our machine learning techniques. First, we define the total length and the principal direction of SFs as:

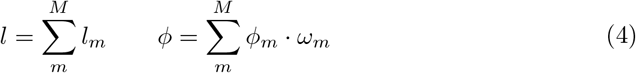

where *l*_*m*_ is the length of *m*-th single stress fiber, *M* is the total number of the stress fibers in an individual cell, *ϕ*_*m*_ is the angle of *m*-th single stress fiber, *ω*_*m*_ = *l*_*m*_| *l* is the weight function. To evaluate each *l*_*m*_ and *ϕ*_*m*_ in every segmented SFs that is image processed by CNN, we apply an edge detection method LSD (Line Segment Detector) [44]. This method allows us to extract straight lines with corresponding two endpoints from the images, and can be used to compute the length and the direction of each SFs. The accuracy of the LSD method compared with the manual extraction is shown in Supplementary Fig. S1.

First, we explore the relationship between the principal orientations of cells and SFs, as well as the relationship between cell area and the total length of SFs. It has been well known that the orientation of stress fibers is closely related to the cell direction [45, 46], and Fig 5A demonstrates the cell direction *ϕ*_*c*_ and the SFs direction *ϕ*_*s*_ are indeed parallel most of the time but still remains a slight disparity. Note that the cell direction *ϕ*_*c*_ is obtained by calculating the angle between the *x*−axis and the major axis of the ellipse that has the same second moments as the cell outline region. Fig 5B shows that the length of stress fibers increases with cell area, which is in agreement with the previous works [47, 48]. To the best of the authors’ knowledge, this is the first report that shows a linear relationship between the length of SFs and the cell area, and this relationship can be fitted with a linear function with a slope 0.172

**Fig 5.**
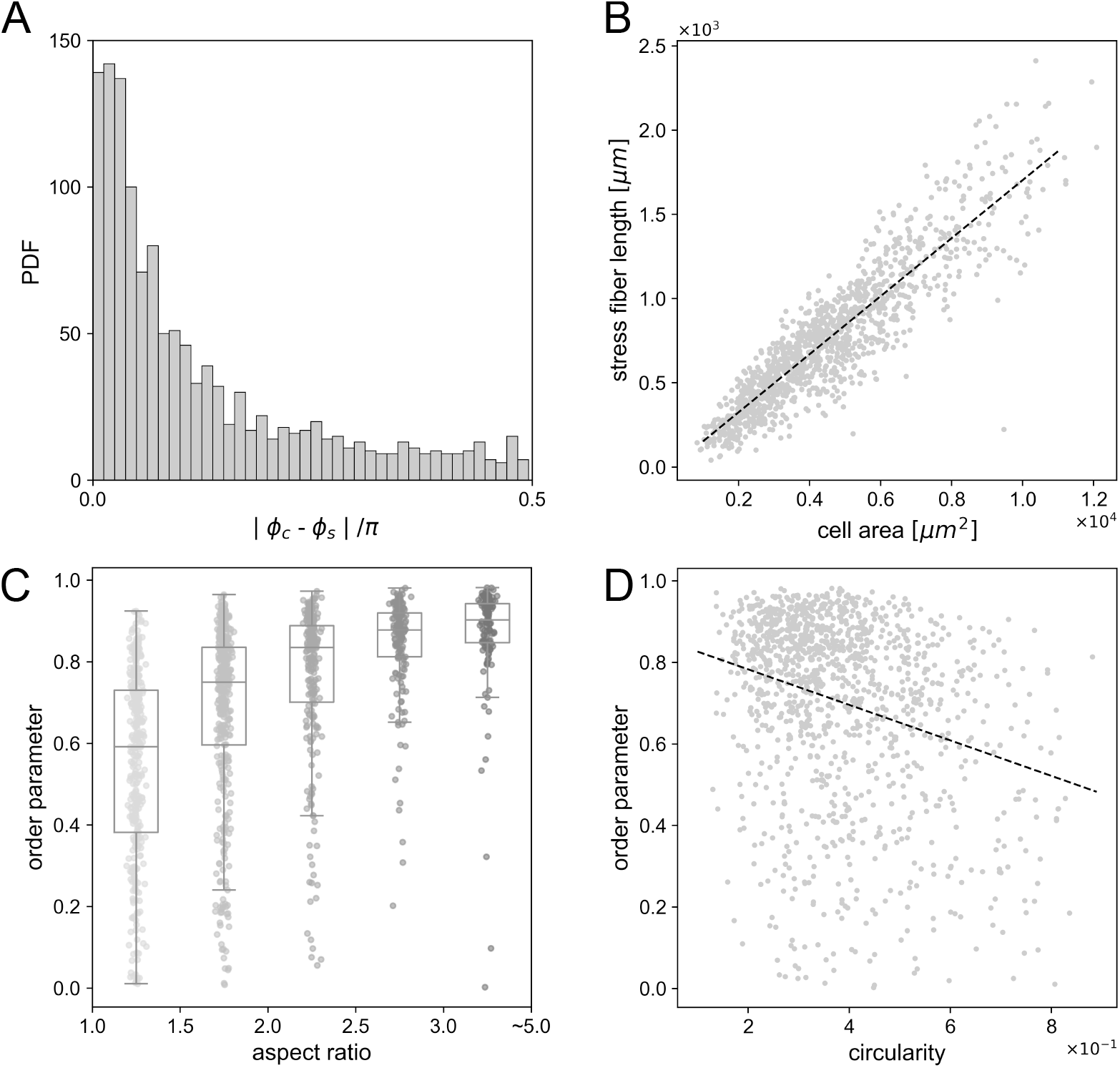
Quantitative analysis of the relation between SFs dynamics and cell morphology. (A) Probability distribution function of the angle differences between cell direction *ϕ*_*c*_ and SFs direction *ϕ*_*s*_. (B) Relationship between SFs length and cell area. (C) Correlation between segmented SFs order parameter *S* and cell aspect ratio, divided into 5 intervals. (D) Correlation between cell circularity and SFs order parameter.

(*ℓ* = 0.172*A*− 0.020 × 10^3^, where *A* is the cell area). Note that this linear relationship *ℓ*∝ *A* can be understood with the following toy model: imagine a rectangular cell shape that has a size *L* × *W* (*L > W*) and an area *A* = *LW*. All the SFs are assumed to have length of *L* and are aligned in the major axis of the cell. When the cell has *n* [1/m] density of SFs in the minor axis, the total length of SF

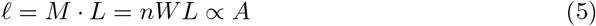

is proportional to the cell area, where *M* = *nW* is the number of SFs.

Second, we focus on how cell morphology interacts with the alignment of SFs. We choose the features of aspect ratio, area, and circularity to compare with SFs alignment. We use the order parameter *S* [49] to quantify this alignment as

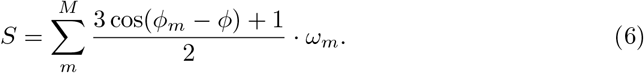

Here, we evenly divided all the SFs data into 5 groups according to the cell aspect ratio from 1.0 to 5.0. As shown in Fig 5C, the average value of order parameter *S* for each interval are 0.548, 0.736, 0.806, 0.869 and 0.899, respectively. The current result is in agreement with previous studies that have shown that the alignment of SFs tends to elongate as the aspect ratio of the micro-patterns that constrain the cell shape [17, 18, 20], or the alignment of actin filaments in microchambers [19]. This illustrates that elongating cells lead to a transition to the longitudinal alignment of the SFs. Meanwhile, as the aspect ratio increases, the IQR (interquartile range) range of the order parameter *S* tends to decrease, demonstrating that the alignments of SFs in a cluster of cells tend to converge as the cell aspect ratio increases. On the other hand, we found no significant correlation between cell area and order parameter *S* (*R* =− 0.032; data not shown), and the cell circularity has a weak correlation with *S* (*R* =− 0.272) as shown in Fig 5D, which means the order parameter decreases with the circularity increases.To summarize the results, the topology of SFs is highly correlated with the low-dimensional expression of cell morphology, especially with the cell aspect ratio. Therefore, we can expect that the localization and alignment of SFs could be predicted from the cell shape.

### Prediction of SFs localization using diffusion model

We now employ our conditional diffusion model to generate SFs from the input cell outline images and evaluate the performance of this approach. Since the results of the diffusion model depend on both input cell shape and Gaussian noise, we generate 100 SFs images from one input cell outline and calculate the average distribution field of all the generated 100 SFs images. We use *P* ∈ [0, 1] to quantify the probability of SFs occurring at a given location, and define the region of *P >* 0.2 as the possible region where SFs are likely to be generated. Visual comparison between the generated SFs results and the actual SFs images are shown in Fig 6A. To describe the quantitative differences, we first define the percentage of intersection region between the ground truth SFs and predicted SF as *S*_⋂_ | *S*_real_, where *S*_⋂_ denotes the overlapped area and *S*_real_ denotes the area of the ground truth. Fig 6B shows that the percentage is 79.3 ± 12.4 [%], which means most of the ground truth SFs are located in the possible region predicted by our diffusion model.

**Fig 6.**
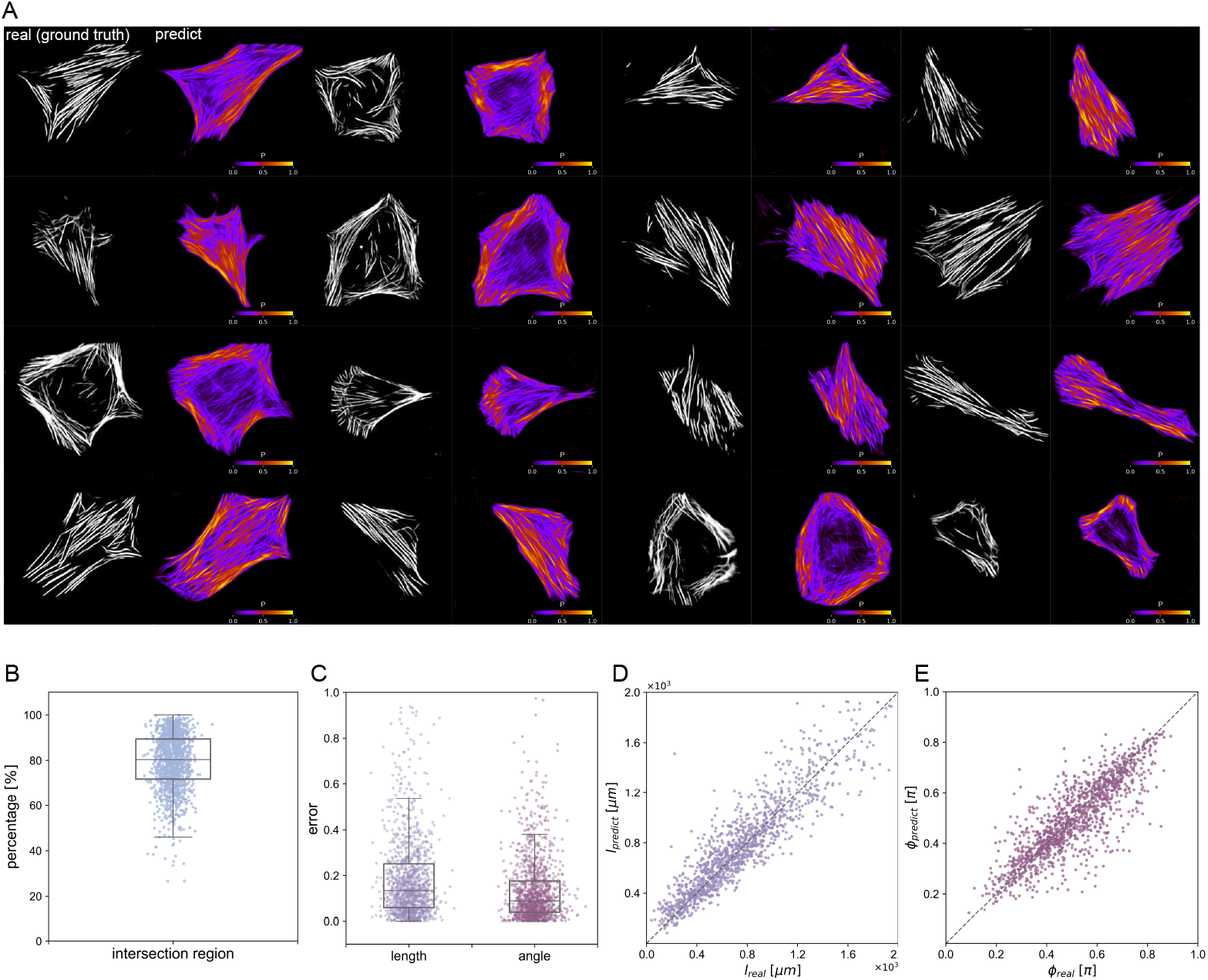
Predicted SFs based on the diffusion model. (A) Comparison of real (ground truth) SFs images and predicted SFs region from the diffusion model. The contour *P* shows the probability of the SFs localization. (B) Percentage of intersection between ground truth SFs and predicted SFs. (C) Error of predicted SFs length *ℓ*^predict^ and direction *ϕ*^predict^ from the diffusion model. (D) Comparison of ground truth SFs length *ℓ*^real^ and predicted SFs length *l*^predict^, with the dashed line indicating equal values. (E) Comparison of ground truth SFs direction *ϕ*^real^ and predicted SFs direction *ϕ*^predict^, with the dashed line indicating equal values.

Next, we compare the generated 1293 cells with the ground truth by measuring the total length *ℓ* and the principal direction *ϕ*. The length and angle are calculated as Eq (4), and the error is defined as *ℓ*^predict^ − *ℓ*^real^ | *ℓ*^real^ and *ϕ*^predict^− *ϕ*^real^ | *ϕ*^real^ where superscripts predict and real indicate the value of the prediction or the ground truth. Note that *ℓ*^predict^ and *ϕ*^predict^ are obtained from the average *P* of the generated 100 SFs images. As shown in Fig 8C, the error of the predicted SFs length is 21.1 ± 31.9 [%], and the principal direction is 13.6 ± 15.2 [%]. Also, Fig D and E show that the length *ℓ* and the principal direction *ϕ* are highly correlated with those of the ground truth.

Additionally, we explored whether the correlation between the generated SFs and the cell morphology remained the same as the actual data. As shown in Fig 7A, the bias between cell principal direction *ϕ*_*c*_ and SFs principal direction *ϕ*_*s*_ can also be noticed by our diffusion model, and the PDF of this bias has a similar distribution with the real data. In Fig 7B, the linear relationship between cell area and SFs length in the predicted data (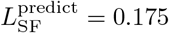 · *S*_cell_ −0.012 × 10^3^) has good agreement with the real cells. The average value of order parameter *S* in Fig 7C for each interval is 0.548, 0.720, 0.804, 0.868 and 0.899, respectively, which also suggests the same results as described in the previous section. Subsequently, the order parameter from the predicted SFs data also correlates with the cell circularity (*R* = −0.242) as shown in Fig 7D. Finally, we find no correlation between cell area and predicted SFs order parameter (*R* = 0.020), which agrees with the real SFs data.

**Fig 7.**
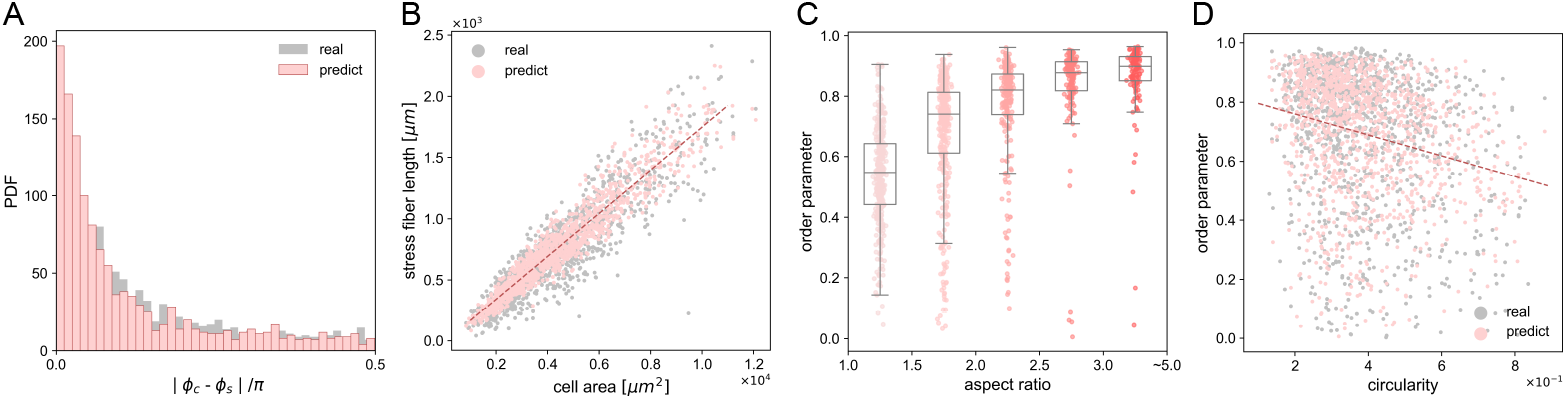
Quantitative analysis of the relation between predicted SFs dynamics and cell morphologies. (A) Probability distribution function of the angle differences between cell direction *ϕ*_*c*_ and SFs direction *ϕ*_*s*_ in both ground truth data and predicted data from the diffusion model. (B) Relationship between SFs length and cell area in both ground truth data and predicted data. (C) Correlation between ground truth SFs order parameter *S* and predicted SFs order parameter *S* from the diffusion model, and cell aspect ratio, divided into 5 intervals. (D) Correlation between cell circularity and SFs order parameter in both ground truth data and predicted data.

As demonstrated above, our diffusion model successfully predicted the localization and distributions of actin stress fibers from the input cell shape images with limited errors. Supplementary S1 and S2 Video further show the application of the proposed system in providing high throughout, real-time prediction of the stress fiber localization during dynamic cell locomotion.

### Virtual experiment: from cell shape to SFs

In previous sections, we successfully predicted the SF localization from the cell shapes using a diffusion model-based machine learning system and evaluated the accuracy. Seeing the current system from a different perspective, it can be said that our system now “installed” the geometric relation between the cell shape and SF localization based on the big data from the experiments; in other words, the system could be used to generate the SF geometry not only from the real cells but even from artificial cell shapes. Although one wants to extract principles from two different parameters, such as the aspect ratio of cells vs SF localizations, it is not trivial to observe cells under ideal situations in experiments. In this section, we show that this system can be utilized to simulate the SF localization for idealized virtual cell shapes. We term this SF generation based on the virtual shape as a “virtual experiment”, and show that this system would be a tool to predict hidden principals of the cell geometry.

First, we generate SFs in a constrained region which has an analogy with the experimental conditions in a previous study [19]. A rectangular-shaped region in this virtual experiment is set with the aspect ratio from 1.0 to 5.0 with a limited area of 2.50 ×10^3^*µ*m^2^. We generate 100 SF images and compute the average SF distribution as visualized in Fig 8A. The order parameter for each constraint is shown in Fig 8B, which shows a continuous increase. The previous study [19] confined the actin filaments into several rectangular microchambers that have a length of 100*µ*m and different aspect ratios by changing width (area). The order parameter from this experiment is shown in Fig 8B as ▽. The increasing trend of the order parameter from the previous study and our simulated results are approximately the same, which demonstrates the potential of our virtual experiment approach.

**Fig 8.**
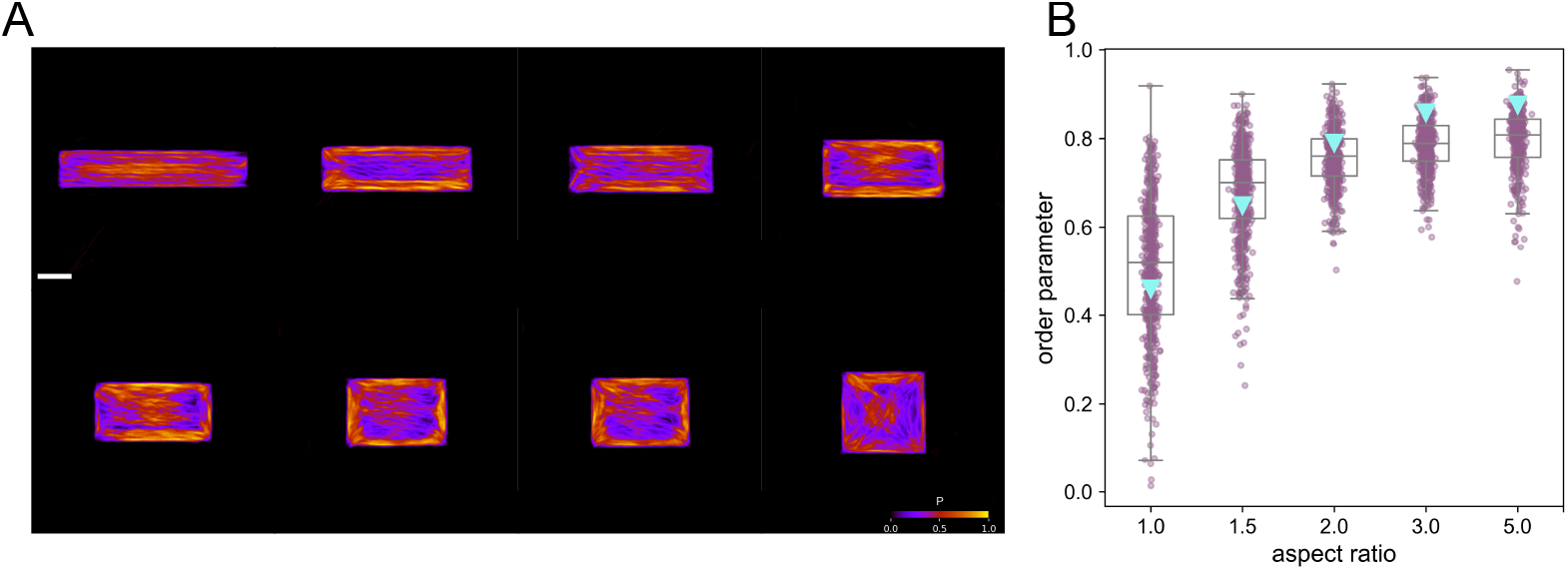
Results from generated SFs comparing with previous studies. (A) Probability *P* of SFs under rectangular constraints predicted by the diffusion model, and a scale bar is 20 *µ*m. (B) Relationship between simulated SFs order parameter and constraint area aspect ratio, compared with results from a previous study (marked as ▽) [19].

Second, we attempt to quantify the geometrical relationship between SFs and the corresponding virtual morphologies. We prepare elliptical shapes with aspect ratios *AR* varying continuously from *AR* = 1 to 5 as the input virtual shape, while the area is set constant 1.96 × 10^3^*µ*m^2^. The generated SFs and the SFs from our experiments are shown in Fig 9A. In the figure, we show generated SFs for two different aspect ratio (*AR* = 1 and 3), and two cells from our experiments that have relatively the same shapes (*AR* = 1.1 and 2.8). Fig 9B shows the average probability *P*_mean_ at the position *r*\*, where *r*\* = *r/R*_max_ is the normalized distance from the cell edge and *R*_max_ denotes the length of the semi-minor axis of the ellipse. As also seen in Fig 9A, the circular cell has a high probability *P*_mean_ ∼ 0.7 at the cell edge while the value is small *P*_mean_ ∼ 0.2 at the center. The profile of probability *P*_mean_ becomes flat for cells with larger AR, and this indicates that SFs homogeneously distribute inside the cells for high AR. Additionally, we visualize the PDF of the probability *P* as shown in Fig 9C (*AR* = 1) and D (*AR* = 3). The PDF of the probability *P* has a single peak at *P* ∼ 0.5 for elongated cell *AR* = 3. On the other hand, the circular cell has a bimodal profile of *P* and the result again suggests that there is a localization of high/low *P* distribution. In order to quantify this trend, we finally show the standard deviation *σ*_*P*_ of the probability *P* in Fig 9E. While the value *σ*_*P*_ is large for small AR since there are bimodalities in the profile, it gradually decreases with AR since the probability *P* becomes more homogeneous inside the cell. Fig 9E shows that cells with *AR <* 2 would show strong localization in the SF distribution.

**Fig 9.**
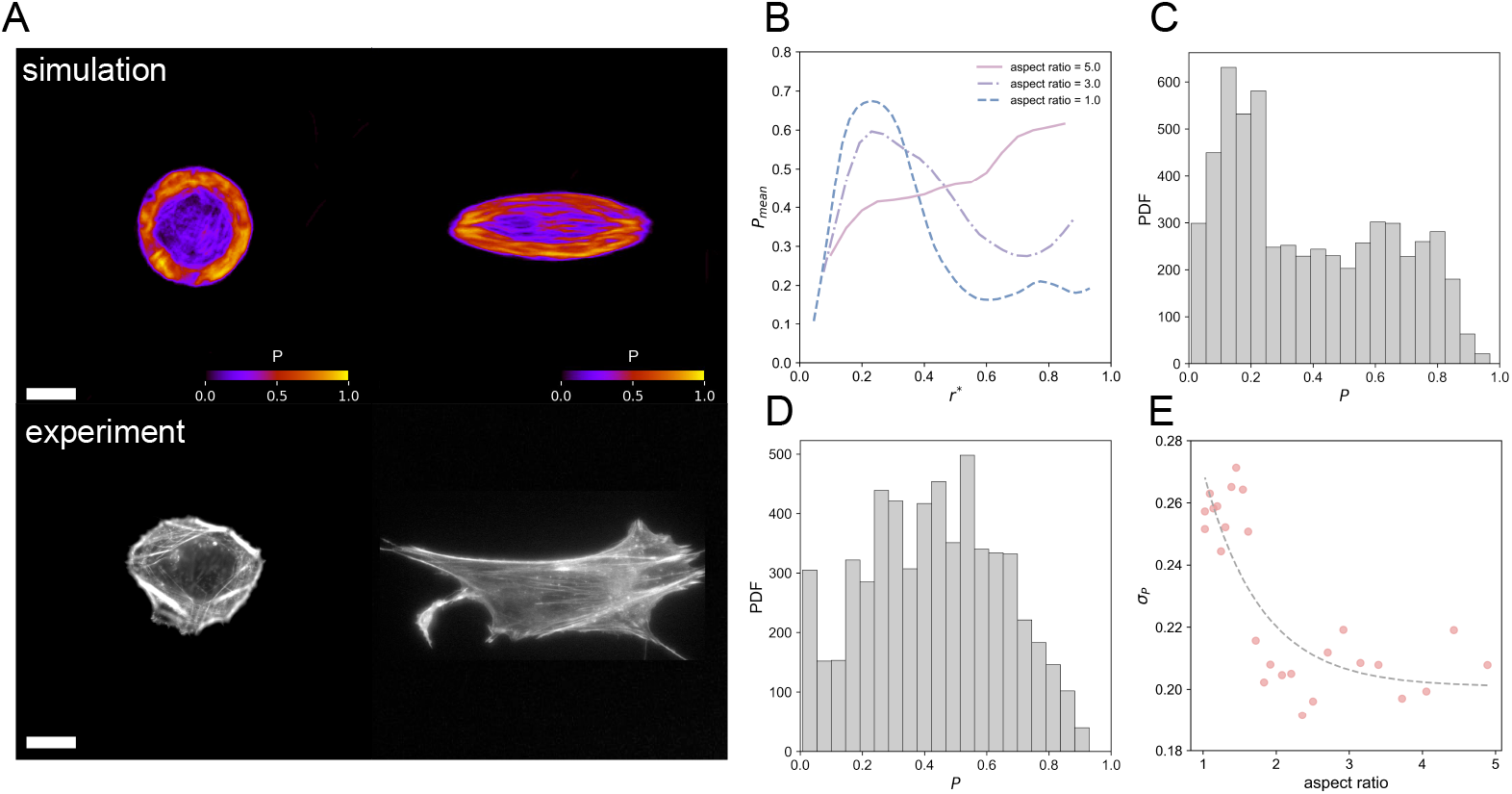
Results from virtual experiment using diffusion model. (A) Comparison of generated SFs with geometrical constraints of aspect ratio (*AR* = 1 and 3) to experimental results of HFF-1 cells with comparable geometry constraints. The cells in the experimental group have aspect ratios of 1.1 and 2.8, and areas of 1.83 ×10^3^*µ*m^2^ and 1.88 × 10^3^*µ*m^2^, which are similar to the morphology conditions during the virtual experiments. Scale bars represent 20 *µ*m. (B) Relationship between distance from the geometrically constrained edge *r*\* and probability of generating SFs *P*_*mean*_ for three cases of aspect ratio (*AR* = 1, 3 and 5). (C,D) Probability distribution function (PDF) of the probability of generating SFs *P* with a constraint of (C) *AR* = 1 and (D) *AR* = 3. (E) Standard deviation *σ*_*P*_ of *P* decreases as constraint aspect ratio increases.

In this final section, we performed virtual experiments using the knowledge “installed” conversion principles that are extracted from our experiments. The generated SFs from the virtual cell shape have good agreement with the previous study of actin filaments. As summarized in the introduction, it is important to understand the geometric relation between cell shapes and cytoskeletons. This conversion, which is based on the big data of experiments, will be a powerful tool to find hidden trends that underly between them as the system can handle and analyze the geometric quantities. For instance, we found that SFs tend to locate at the cell edge for *AR <* 2 while the probability *P* is expected to be more homogeneous for cells with large aspect ratios. This concept of “installing” the underlying rules can be used not only for this specific task, but can also be extended to other conversions to simulate the characters of cell nature.

## Conclusion

We propose a diffusion model-based machine learning system, which allows for predicting the geometries and localizations of SFs from cell morphologies. First, we extract SFs and cell shapes from raw fluorescence microscopic images using CNN and image processing methods. From the extracted SFs, we found the total length of SFs in a single cell linearly increases with the cell area. We also showed that the order parameter of SFs increases with the cell aspect ratio. Note that we managed to draw these conclusions by processing big data with the help of machine learning. Second, we train the system with the diffusion model using corresponding two image sets, namely SFs images and cell shape images, as training data. The predicted SFs are highly correlated with our experimental data, displaying an overlap area of 79.3 ± 12.4 [%] with the actual SFs observed in the experiments. The errors were 21.1 ± 31.9 [%] for SFs length and 13.6 ± 15.2 [%] for SFs principal direction. Finally, we show that our system can be utilized for the virtual experiments to predict the localization of SFs from artificial cell shapes as the inputs.

Cell biology has long examined the connection between cellular morphology and function [50] and has studied individual cells in detail [9, 11, 51]. The ability of cells to contract, which is largely driven by actin SFs, has been linked to a range of processes such as proliferation, differentiation, apoptosis, and tumorigenesis. Cell shape is one of the most common cellular features and can be obtained relatively easily through optical microscopes and, in the meantime, actin SFs are vital components of cells in controlling many cellular functions, with effects ranging from migration and proliferation to apoptosis. Considering the complexity of the cellular biological mechanisms along with the physical/biological process associated with SFs dynamics and the limitation of the throughput in experiments, we believe this “installable” knowledge of conversion from cell geometry to SFs could be a powerful system, since this approach provides a virtual environment that can be applied to quantitative analysis on the correlations between the cell morphologies and the corresponding actin distribution characteristics. By combining with our previous study [52, 53], which is a machine learning based approach that can extract cellular force distributions from microscope images, we are planning to analyze how cell morphologies are connected with the cell mechanics (see supplementary S3). Our approach provides a powerful and cascadable framework to understand the geometric features of cells, which would support new findings in the field of cell biology and mechanobiology.

## Supporting information

**Fig S1.**
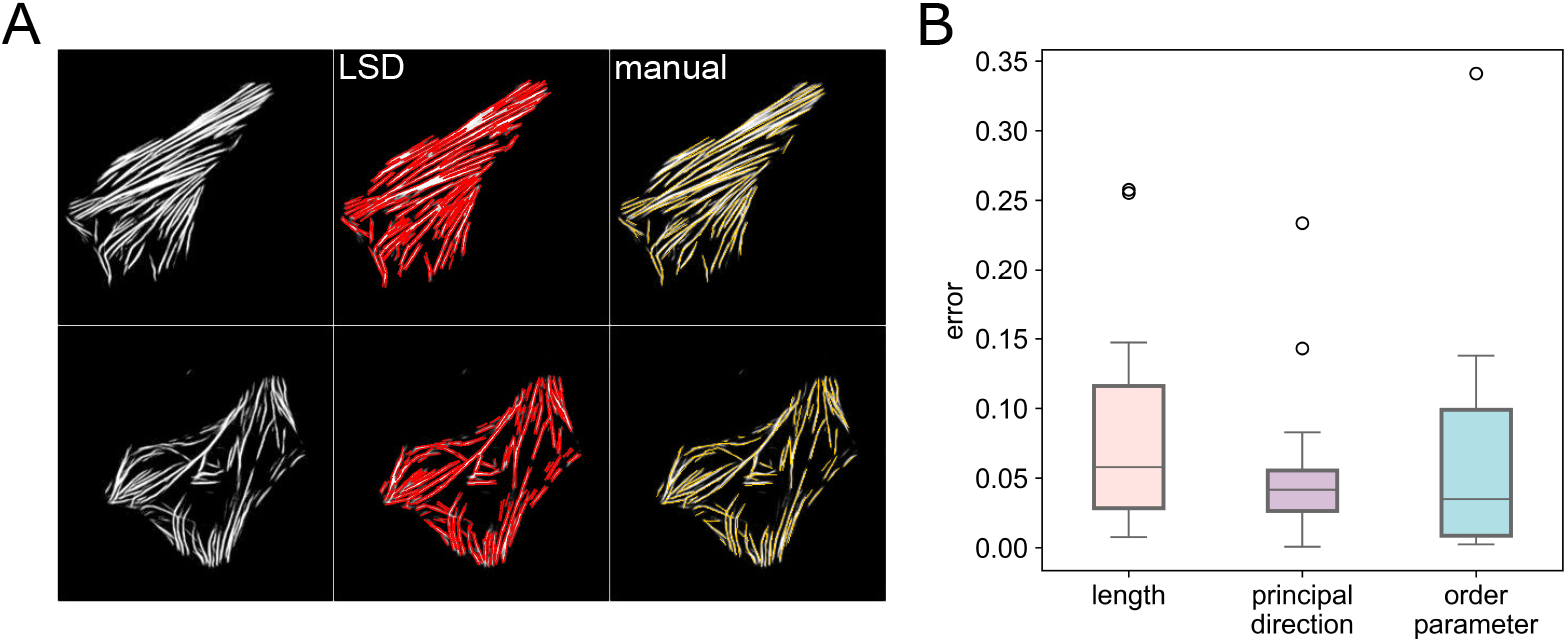
Errors of LSD method. Due to the large amount of acquired SFs experimental data and predicted data from the diffusion model, we use the LSD method to automatically determine the total length, principal direction, and order parameter of the SFs in a single cell. Figure A shows the results detected by the LSD method and those labeled manually. Figure B summarizes the error between the LSD method and the manual approach (*N*_test_ = 20). The errors in SFs length, principal direction, and order parameter are 7.12 ± 8.50 [%], 5.28 ± 5.11 [%], and 6.81 ± 7.79 [%], respectively. These results demonstrate that the LSD method is effective for automatically analyzing the characteristics of SFs extracted from CNN or generated by the diffusion model.

**S1 Video. Prediction of SFs from living microscopic image (i)**. Extract HFF-1 cell shapes from live videos and generate corresponding SFs data using a trained diffusion model.

**S2 Video. Prediction of SFs from living microscopic image (ii)**. Same as S1 Video.

**S3 Video. Combining predicted SFs data with WFM method**. Combining with our previous study WFM (Wrinkle Force Microscopy), we believe for the future work, finding cell mechanism features only from cell morphology could be possible.

## Acknowledgments

This work is supported by JSPS KAKENHI Grant Number 21J22170, Chinese Scholarship Council (CSC) Scholarship.

